# networkGWAS: A network-based approach to discover genetic associations

**DOI:** 10.1101/2021.11.11.468206

**Authors:** Giulia Muzio, Leslie O’Bray, Laetitia Meng-Papaxanthos, Juliane Klatt, Karsten Borgwardt

## Abstract

While the search for associations between genetic markers and complex traits has led to the discovery of tens of thousands of trait-related genetic variants, the vast majority of these only explain a small fraction of observed phenotypic variation. One possible strategy to detect stronger associations is to aggregate the effects of several genetic markers and to test entire genes, pathways or (sub)networks of genes for association to a phenotype. The latter, network-based genome-wide association studies, in particular suffers from a vast search space and an inherent multiple testing problem. As a consequence, current approaches are either based on greedy feature selection, thereby risking that they miss relevant associations, or neglect doing a multiple testing correction, which can lead to an abundance of false positive findings.

To address the shortcomings of current approaches of network-based genome-wide association studies, we propose `networkGWAS`, a computationally efficient and statistically sound approach to network-based genome-wide association studies using mixed models and neighborhood aggregation. It allows for population structure correction and for well-calibrated *p*-values, which are obtained through circular and degree-preserving network permutation schemes. `networkGWAS` successfully detects known associations on semi-simulated common variants from *A. thaliana* and on simulated rare variants from *H. sapiens*, as well as neighborhoods of genes involved in stress-related biological processes on a stress-induced phenotype from *S. cerevisiae*. It thereby enables the systematic combination of gene-based genome-wide association studies with biological network information.

**Availability:** https://github.com/BorgwardtLab/networkGWAS.git

## 1 Introduction

Genome-wide association studies (GWAS) aim to identify statistical associations between genetic variants– most commonly in the form of single nucleotide polymorphisms (SNPs)–and disease risk or other phenotypes. However, most of the phenotypes of interest are complex traits in the sense that they do not follow a Mendelian pattern of inheritance since they are controlled by multiple SNPs and genes, and are influenced by environmental factors. Traditional GWAS face the fundamental obstacle of *missing heritability* with respect to such traits, i.e., that single SNPs which were found to be significantly associated often account for a small portion of the variation of heritable phenotypes. As previously argued, large parts of missing heritability could be due to genetic interactions–if the development of a certain phenotype involves interaction among multiple pathways–rather than directly correspond to undetected association with genetic variants [31]. Therefore, a great effort has been undertaken to develop more comprehensive and powerful GWAS methodologies, aiming at understanding and incorporating biological mechanisms underlying the genetics of complex traits. To date, the rich knowledge about biological networks which is already available–such as protein-protein interaction (PPI) and gene regulatory networks–is rarely leveraged in a statistical and rigorous way in GWAS. Including such contextual and functional information, representing processes relevant to the phenotype under study, can enable an increase in statistical power as well as improve interpretability in GWAS aimed at complex traits, thus representing a promising approach to overcome the problem of missing heritability.

The problem of limited power in GWAS is generally rooted in both a large marker-to-sample ratio and low heritability of complex traits. In order to mitigate that, two strategies have been pursued: (i) to group genetic markers and test them at once, thereby reducing the multiplicity of markers tested [9, 14, 17, 23], or (ii) to employ biological networks in order to conduct a *post hoc* aggregation of association [1, 2, 6, 8, 10, 25, 29]. Both approaches amplify the signal of SNPs or genes which are collectively phenotype-related but would not pass the significance threshold on their own. However, within the set-based test strategy, so far, SNP sets are typically chosen based on membership to a functional unit on the genome. Hence, this strategy lacks a principled procedure to select SNP sets that goes beyond single genes or mere regions on the genome. The *post hoc* aggregation strategy, on the other hand, suffers from the absence of statistically sound *p*-values for the set of aggregated SNPs. We propose to combine both strategies and thereby overcome their respective weaknesses. More precisely, our approach entails testing sets of SNPs, as done for example by the FaST-LMM-Set method [17], but we guide the SNP selection by means of biological networks. Thus, we arrive at a strategy that incorporates both a biologically meaningful way to select SNP sets that goes beyond functional units, and that yields statistically rigorous *p*-values for the SNP sets tested.

## 2 networkGWAS

In this section, we detail how we exploit the biological network information, we discuss the mathematical model we use and the details of how we obtain *p*-values.

### 2.1 Neighborhood aggregation

We test pre-defined sets of SNPs, rather than single SNPs, in order to both reduce the number of markers tested and to account for gene interaction in addition to mere genetic variance. Multiple methods performing SNP set-based tests already exist (including gene enrichment analysis [9], collapsing methods [14], multivariate regression [23], and linear mixed models (LMMs) [17]). However, none of them incorporates biological network structure in order to guide the SNP-set selection, thereby choosing SNP sets that are not representative of biological mechanisms. In our approach, instead, we select SNP sets to be tested based on protein-protein interaction (PPI) networks, i.e., the graph representation of the interactions between proteins. PPIs are essential for almost all biological mechanisms, and are defined as the specific, non-generic, physical contact between proteins in a particular biological context [22]. These interactions can be both stable (e.g., as in multi-enzyme complexes) or transient (e.g., as in interaction with kinases [11]). Since we focus on complex phenotypes, we employ the entire PPI network–including both stable and transient interactions–thus capturing effects of molecular mechanisms taking place in diverse cells and tissues, and of various kinetics.

More precisely, each sample *i* in the GWAS dataset is represented as a graph *G_i_* = (*V, E*), where *V* is the set of nodes and *E* is the set of edges in the PPI network. In these graphs, the nodes *V* represent genes, and edges *E* indicate any kind of PPI between gene products of the two nodes they connect. Since each sample uses the same PPI network, the topology is shared, i.e., (*V, E*) is the same for each sample. The node labels, however, vary depending on the sample. Each node *v* ∈ *V* is attributed with a feature vector 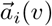 comprising the values of all SNPs overlapping with the corresponding gene. Based on this representation, one SNP set per gene is constructed by means of concatenating the feature vector of the gene itself as well as its *k*-hop neighbor genes according to the PPI network. As a result, the node label vector 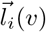 of a node *v* and a sample *i* is now represented by the union of its own SNPs and those from its *k*-hop neighborhood 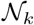,

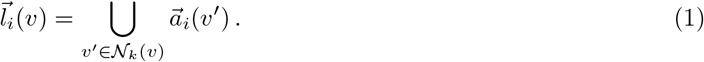

This neighborhood aggregation operation is visualized in panel (a) of Fig. 1. We thereby directly test the significance of biological subnetworks to identify pathways underlying complex phenotypes. In summary, our neighborhood aggregation approach is akin to the idea underlying graph kernels [4] or graph convolutional networks (GCNs) [12]. All of these methods leverage localized first-order approximations of subgraph structure in order to avoid an exhaustive search of all subgraphs, which would scale exponentially in the size of the network at hand, whereas our approach is linear in the number of nodes, even when the *k*-hop neighborhood is defined to be greater than 1. Note that in all the experiments we perform, we employ 1-hop neighborhoods to define our SNP sets.

**Fig. 1:**
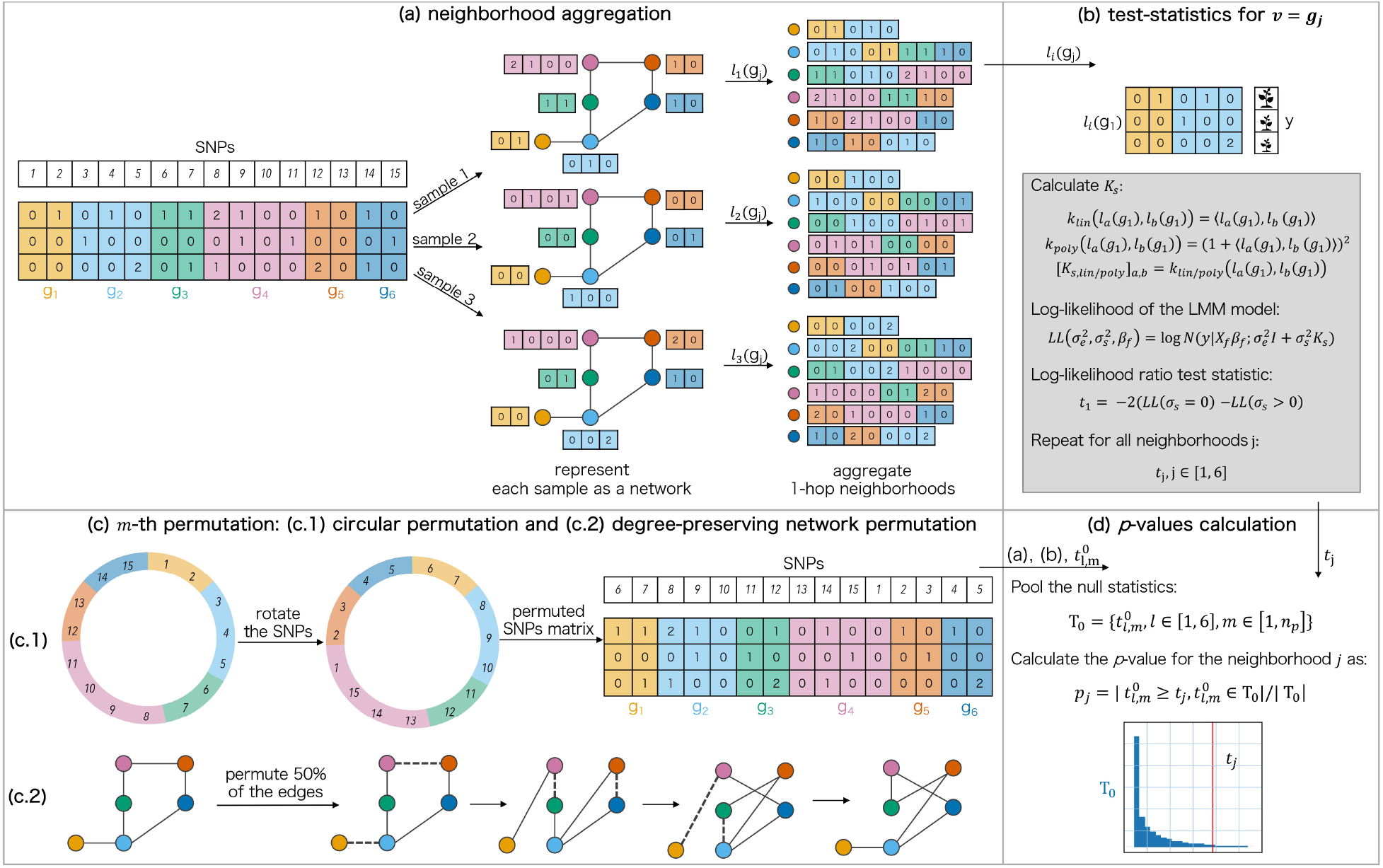
Overview of `networkGWAS`: (a) given a SNP matrix, a PPI network, and a mapping of SNPs onto the genes *g_j_* (color-coded), the first step, i.e., the neighborhood aggregation, begins by representing each of the samples as a network which shares a fixed topology from the PPI network but differs in the node feature values. Subsequently, the 1-hop neighborhood aggregation of the features is performed for each sample *i*, resulting in each node *j* being labeled with *l_i_*(*g_j_*)–the concatenation of its own features and the features of its 1-hop neighbors in the PPI network. (b) Per each gene *j*, either *K*_lin_ or *K*_poly_ is calculated from *l_i_*(*g_j_*), ∀*i* ∈ [1, 3]. Then, the values of Eq. (2) for *σ_s_* = 0 and *σ_s_* > 0 are estimated via restricted maximum likelihood, and, subsequently, the likelihood-ratio test statistics *t_j_*, *j* ∈ [1, 6] are obtained. (c) The distribution of the test statistics under the null hypothesis of no association signal between the neighborhoods and the phenotype is derived via a permutation procedure which combines a circular permutation (c.1) with a degree-preserving network permutation (c.2). Having obtained the permuted settings, steps (a) and (b) are performed to obtain a test statistic 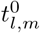 per neighborhood *l* and permutation *m*. (d) 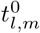, ∀*l* ∈ [1, 6] and ∀*m* ∈ [1, *n*_p_] are pooled to obtain the null distribution 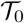. Afterwards, a *p*-value per each neighborhood is estimated by calculating the ratio between the number of null statistics 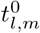 that are greater than or equal to the *t_j_*, as obtained on the non-permuted setting, divided by the total number of 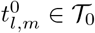.

### 2.2 Model

Once SNP sets have been selected in the aforementioned manner, we employ a FaST-LMM-Set like model [16] to estimate the statistical associations with the phenotype of choice. The LMM we use,

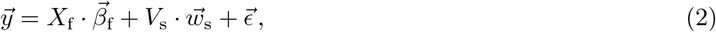

features one random effect 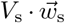, which accounts for the similarity among the SNPs of the set to be tested. The precise form of *V_s_* depends on the similarity measure chosen and will be specified by means of Eqs. (4) to (6) below. Above, *y* contains the continuous phenotype values of the *n* individuals studied, *X_f_* is the *n* × *n*_f_ matrix of *n*_f_ fixed effects (e.g., a column of 1s corresponding to the intercept and other covariates, see Section S.1; here and throughout the manuscript, all section, figure, and table numbers starting with an “S” refer to the Supplementary), 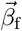 denotes the vector of the *n*_f_ fixed effect weights, 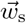 comprises the signal, i.e., the random effects of the *n*_s_ SNPs of interest and included in the pre-defined SNP set to be tested, and 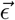 models residual noise. 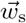, and 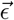 are assumed to be drawn from multivariate Gaussian distributions, respectively 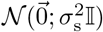 and 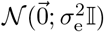, where 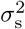 represents the genetic variance, 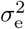 the residual variance, and 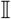 the identity matrix. Marginalizing over fixed effects, the log-likelihood of the model (2) reads

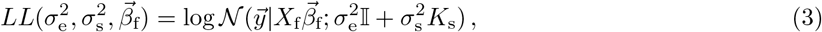

where *K_s_* is the covariance matrix capturing the similarities among the SNPs in the test set. The parameter *σ_s_* serves to distinguish the null model (i.e., *σ_s_* = 0) from alternative models (i.e., *σ_s_* > 0), and is estimated from the GWAS dataset by means of restricted maximum likelihood. While in the original FaST-LMM-Set, *K*_s_ measures similarity through a linear kernel *k*^(lin)^, we additionally explore the use of a quadratic kernel *k*^(poly)^ for *K*_s_ in our method:

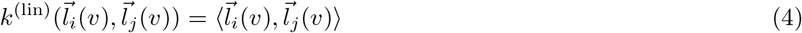

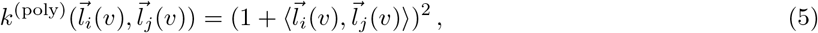

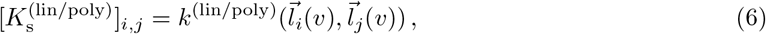

where 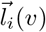 and 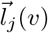 are defined according to Eq. (1) for the *i*-th and the *j*-th sample. We chose an inhomogeneous polynomial kernel in order for it to be able to capture both linear and non-linear similarity. In an additional deviation from FaST-LMM-Set, we normalize the diagonal entries of our final kernel matrix 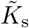 to be 1, by means of the following equation:

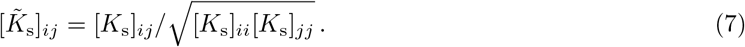

Note that the definition of *V*_s_ in Eq. (2) varies depending on the type of the kernel we use. If the kernel is defined following Eq. (4), e.g. the linear kernel, *V*_s_ corresponds to the *n* × *n*_s_ design matrix whose rows correspond to *l_i_*(*v*), *i* = 1,…,*n*, normalized to be consistent with Eq. (7). When the kernel is instead calculated according to Eq. (5), *V*_s_ constitutes a higher-dimensional vector in the reproducing kernel Hilbert space which additionally accounts for the *n* × *n*_s_(*n*_s_ – 1) second-order interactions among the SNPs in the test set.

### 2.3 *p*-value computation

`networkGWAS` aims to identify neighborhoods that are statistically associated with a trait of interest. In the following, we refer to the *k*-hop neighborhood of gene *v_j_* as 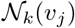, with *j* = 1,…, *n*_g_ and *n*_g_ equal to the total number of genes. We then define a null hypothesis *H_j_* for each neighborhood, which signifies that 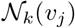 does not affect the phenotype. Explicitly, this states that the set of SNPs representing 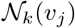, i.e., *l_i_*(*v*), *i* = 1,…, *n*, does not exhibit an association signal with the trait under study. Note that under the null hypothesis *H_j_*, the patterns of linkage disequilibrium (LD) among the SNPs and the population structure, if present, must be preserved. LD patterns represent the non-random correlations between the SNPs due to, e.g., spatial proximity on the genome, while population structure refers to the complex relatedness among the individuals, which impacts the values and structures of both the SNPs and the phenotypes. Having defined *H_j_*, we further need to

1. Define a measure, i.e., a test statistic, to quantify the association signal between each neighborhood and the phenotype,
2. Obtain the null distribution of the test statistics underlying the null hypothesis,
3. Estimate the *p*-values, and
4. Define a strategy to identify the statistically significantly associated neighborhoods.

Since we rely on FaST-LMM-Set for our SNP set-based test, the association between 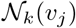 and the phenotype is quantified by calculating the log-likelihood ratio between the maximum restricted likelihood estimate of the alternative and null models from Eq. (3). The test statistic obtained in this way for 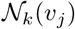 is referred to as *t_j_*. Unlike the original FaST-LMM-Set method which uses a parametric null distribution, we adhere to a non-parametric distribution for `networkGWAS` instead. The reasoning behind this choice is detailed in Section S.3. We determine the distribution of test statistics under the null hypothesis by means of a permutation strategy that allows to destroy the association signal between the sets of SNPs (i.e., the neighborhoods) and the phenotype, while simultaneously preserving LD patterns and population structure. We achieve this by performing permutations on both the SNPs and the network level, namely a circular permutation of the SNPs and a degree-preserving permutation of the network, both of which are detailed in the following.

#### Circular permutation of the SNPs

The implementation of the circular permutation procedure is inspired by Cabrera *et al*. [5]. We consider the SNPs to be ordered according to their genomic position in a circular way, namely that after the last SNP on the last chromosome, one restarts from the first SNP on the first chromosome. A single permutation is then performed by randomly selecting a number between 1 and the total number of SNPs (which in our case would be the total number of SNPs across all the *n*_g_ neighborhoods), and then rotating the SNPs by that value, while keeping the genomic coordinates fixed. A visualization of this is reported in panel (c.1) of Figure 1. This operation causes the SNPs to be assigned to different genomic positions compared to their original location on the genome (therefore positionally mapping them to a different gene compared to non-rotated scenario), while conserving the same position with respect to each other, hence preserving the LD patterns among the SNPs. This operation sufficiently preserves any confounding population structure since the relative position between SNPs is again maintained. Furthermore, the relatedness signal is also preserved when constructing the null distribution since the arrangement of the individuals in the SNPs to test *V*_s_ (or interactions of SNPs to test), phenotype (*y*), and fixed effects (*X*_f_) is not varied. Note that when performing multiple circular permutations, only non-repeating rotations are considered.

#### Degree-preserving network permutation

We permute the network on top of permuting the SNPs to add a level of randomization when performing the neighborhood aggregation operation. This network permutation consists of a methodology that allows us to shuffle the edges while maintaining the degree of each node preserved, thereby enforcing a comparable graph structure despite the permutations. Consider a network *G* = (*V, E*), where *V* is the set of nodes *V* = {*v_j_*, *j* = 1,…, *n*_g_}, and *E* is the set of edges. To generate one permutation of the network, the following steps are performed until 50% of the edges are rearranged:

1. Randomly select two pairs of connected genes, (*v_a_*, *v_b_*) and (*v_c_*, *v_d_*).
2. Check whether *v_a_* or *v_b_* are connected with *v_c_* or *v_d_*; if so, return to point 1, otherwise proceed to point 3.
3. Remove the edge between *v_a_* and *v_b_*, and *v_c_* and *v_d_*.
4. Connect *v_a_* with *v_c_*, and *v_b_* with *v_d_*.

Figure 1(c.2) shows an example of this permutation technique. Note that the only parameter to set for this two-level permutation strategy is the percentage of edges to shuffle, which we set to 50%.

After having permuted the SNPs and the network following the aforementioned circular and degree-preserving permutations respectively, we can again define neighborhoods on these permuted scenarios (as detailed in Section 2.1), and, subsequently, we can calculate a test statistic for each of them. Specifically, we can obtain 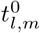 for the *l*-th neighborhood and the m-th permutation as described above. By pooling these test-statistics obtained from all neighborhoods under all permutations, we obtain an empirical test-statistic distribution under the null hypothesis [30]. Note that with this procedure, we obtain *n*_g_ statistics per each permutation, decreasing the total number of permutations *n*_p_ required (discussed in Section S.4).

Having determined tj and the null distribution of these statistics under the null hypothesis, i.e. 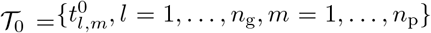, we calculate the *p*-value for 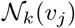 as 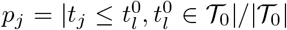. Note that for avoiding significance values of zero in case 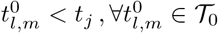, a pseudocount can be added. This procedure allows us to obtain calibrated *p*-values which measure the significance of the statistical association of a particular neighborhood of interacting genes and the phenotype. More precisely, the *p*-values represent a well calibrated distribution and a genomic inflation factor close to 1, i.e., *λ_GC_* ∈ [0.8, 1.2], for 98% of the analyses performed on the common variants scenarios, namely for the *A. thaliana* semi-simulations, and for both *A. thaliana* and *S. cerevisiae* real data. The results of the synthetic simulations on the *H. sapiens* rare variant report a slight deflation in the distribution of the high-range *p*-values (i.e., *p*-values close to 1). Despite this, the low-range *p*-values are well-calibrated, and are supported by the obtained competitive performance, reported in Section S.8.2. For these reasons we retain this use case as well. For further details on the obtained *p*-values, the reader is referred to Section S.3.

In a last step, we identify the neighborhoods that are statistically associated with the phenotype of interest. When studying a particular organism, often multiple related phenotypes are available. Hence, we are testing multiple hypotheses on two levels, namely *n*_g_ neighborhoods for *T* traits. Therefore, to account for this aspect when correcting our significance level for multiple testing, we employ the hierarchical testing procedure proposed by Peterson *et al*. [21] (detailed in Section S.5), which is based on the Benjamini-Hochberg procedure [3] and allows to effectively control the false-discovery rate (FDR) in such scenarios. When performing our analyses, we control the FDR at level 0.05.

Lastly, the computational cost involved are in the order of 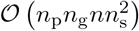. Refer to Section S.2 for the details.

## 3 Simulations

In this section, we detail the simulations studies we perform, designed such to best demonstrate the robustness and limitations of `networkGWAS` under varying conditions, and we introduce the state-of-the-art methods we contrast our results to. In particular, we apply our method and the comparison partners on semi-simulated settings from *A. thaliana* and fully synthetic settings from *H. sapiens*. While the first is presented in the following, the *H. sapiens* use case is discussed in Section S.8.2.

### 3.1 Experimental setup

In order to apply our method, one needs a GWAS dataset consisting of genotypes and a phenotype of interest, as well as a PPI network relevant to the phenotype chosen. In our experiments we employ natural genotypes and a PPI network in combination with simulated phenotypes. This allows us to test our method on genotypes with realistic LD and MAF patterns and on a biological network with sensible structure, while also enabling us to have a ground truth to compare our results to. We use the genotype dataset for *A. thaliana* from the AraGWAS Catalog [28] and the PPI network from The Arabidopsis Information Resource (TAIR) database [13]. More details on the data and processing in Section S.6.1. Note that we chose the TAIR network for our semi-simulated setting since its smaller size allows us to run a large number of fast experiments.

To simulate the phenotypes, we firstly define the following parameters: (a) the number of genes *n*_cg_ that carry causal SNPs, henceforth called causal genes, (b) the ratio of causal SNPs on a causal gene, *r*_c_, (c) the mean ratio of causal neighbors (RCN) of a causal gene, (d) the signal-to-noise-ratio (SNR), and (e) the mixing ratio of linear-to-nonlinear (RLN) signal. Note that the causal genes (SNPs) are selected among the 1,327 genes (37,458 SNPs) that remain after the processing. Therefore, the simulated genotypephenotype associations are modeled within this subset of the genome. All the above defined parameters are systematically varied in our experiments. Thus, we investigate the robustness of our method’s and comparison partners’ performance as we depart from the most amenable scenario (S0) of a purely linear signal, spread across a single or very few causal subgraph(s) composed of a high number of causal genes, with a high ratio of causal neighbors, a realistic signal-to-noise ratio, and a high ratio of causal SNPs on the causal genes. Table S.1 summarizes the simulated scenarios, starting from our anchor scenario S0 and deviating from its conditions by:

– Varying the SNR while keeping the RCN, RLN, *r*_c_, and *n*_cg_ constant,
– Varying the RLN while keeping the SNR, RCN, *r*_c_, and *n*_cg_ constant,
– Varying the RCN while keeping the SNR, RLN, *r*_c_, and *n*_cg_ constant,
– Varying the *r*_c_ while keeping the SNR, RCN, RLN, and *n*_cg_ constant,
– Varying the *n*_cg_ while keeping the SNR, RCN, RLN, and *r*_c_ constant.

We thereby study the isolated effects of moving towards more challenging/different RCN, RLN, SNR, *r*_c_, and *n*_cg_ respectively. For each scenario, we simulate 5 phenotypes as:

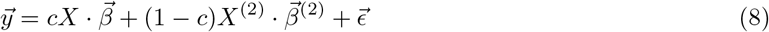

where *X* is the *n* by *p* matrix of all SNPs across the genes considered in the analysis (i.e., 37,458 in this case), 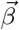 represents the fixed effect of these SNPs, 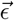 models the noise, *X*^(2)^ is the *n* by *p*(*p* – 1)/2 second-order design matrix of all SNP interactions, 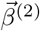 comprises the fixed effects of all SNP interactions, and the coefficient *c* serves to tune between the ratio of linear to non-linear signal. The choice of *c* and 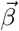 leads to the realization of the different scenarios, as detailed in Section S.7.

### 3.2 Comparison partners and performance evaluation

We compare `networkGWAS`’s performance to: (i) the original FaST-LMM-Set approach [17], however, with sets based on single genes rather than neighborhoods of interacting genes as defined by the PPI network, (ii) NAGA [6], and (iii) dmGWAS [29]. Both NAGA and dmGWAS incorporate PPI information following a *post hoc* strategy. In fact, they both commence with a classical GWAS analysis to obtain single-SNP *p*-values. Subsequently, dmGWAS employs a greedy-selection based, dense module searching, aiming to find PPI subnetworks enriched in low *p*-value SNPs. NAGA, on the other hand, first represents and scores entire genes based on their most significant SNP and then relies on a PPI-network propagation approach in order to spread and revise scores across gene-neighborhoods. While `networkGWAS` and FaST-LMM-Set approach return a *p*-value per each neighborhood or gene, dmGWAS and NAGA provide a score for the subnetworks and genes respectively. Having obtained these *p*-value-based and score-based rankings, we evaluate the performance of linear and non-linear `networkGWAS` as well as the comparison partners by means of their respective mean area under the precision-recall curve (AUPRC) in terms of causal genes. Here, linear and non-linear refers to the SNP-set kernel employed (see Eq. (5)), and the mean refers to the average with respect to the different random realizations of the various simulation settings.

### 3.3 Results

An overview of the results is shown in Figure 2. The top left panel depicts the full mean precision-recall curves for all methods studied under the conditions of our anchor scenario S0, which is most amenable to network guided search for genotypic-phenotypic associations (see Table S.1). In this scenario of a purely linear signal, spread across a single or very few causal subgraph(s) with a high ratio of causal neighbors, and high signal-to-noise ratio, both linear and non-linear `networkGWAS` substantially outperform all comparison partners by achieving an AUPRC of 76.4%±20.5% and 75.9%±20.3%, more than doubling the average AUPRC of the strongest competitor, NAGA, which averagely reports an AUPRC of 28.4%±5.2%. When dividing `networkGWAS`’s AUPRC by the prevalence, 3.8% in this scenario, we obtain a factor of about 20 for both linear and nonlinear `networkGWAS`, further highlighting `networkGWAS`’s strong performance in our anchor scenario. Furthermore, it is worth noting that `networkGWAS` presents higher recall in comparison to dmGWAS per each precision value, highlighting the improvement in the identification of relevant associations achievable by employing neighborhood aggregation instead of greedy search-based algorithm.

**Fig. 2:**
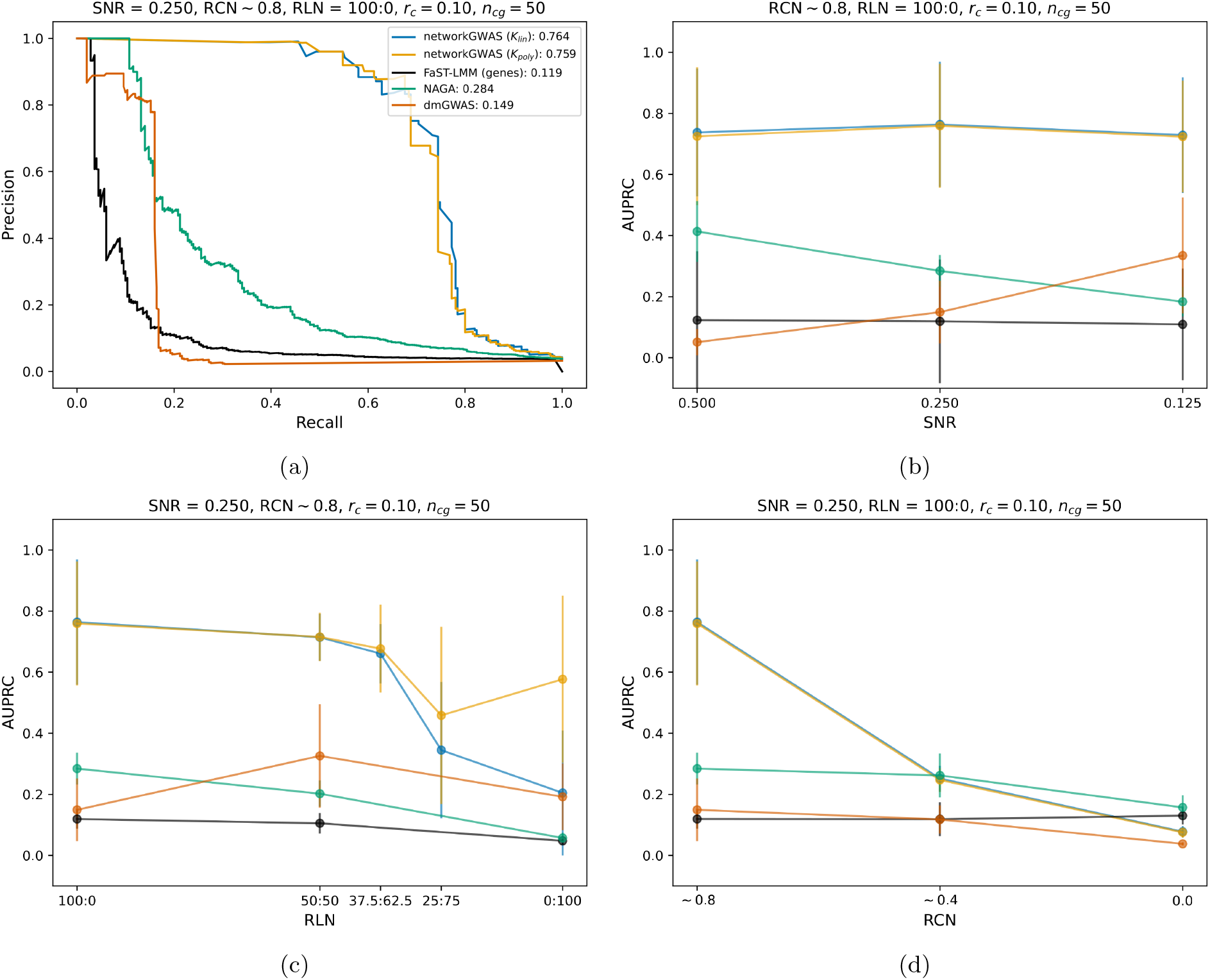
Results from simulating the phenotypes of *A. thaliana*. We present results from our method (`networkGWAS`), using either a linear (*K*_lin_) or a polynomial (*K*_poly_) kernel, as well as the performance of other comparison methods. In Subfigure 2a, we show the AUPRC of the baseline scenario (S0). In Subfigures 2-b-2d, we vary one variable while keeping the other four fixed: the signal-to-noise ratio (SNR) (2b), the mixing ratio of linear-to-nonlinear (RLN) signal (2c), the mean ratio of causal neighbors (RCN) (2d). The AUPRC in function of the ratio of causal SNPs on a causal gene (*r*_c_) and the number of causal genes (*n*_cg_) is shown in the Supplementary.

As demonstrated in the top right panel of Figure (2), this dominance in performance is invariant as one departs from the conditions of S0 by varying the SNR while keeping the purely linear nature of the signal and the ratio of causal neighbors, the percentage of causal SNPs on a causal gene and the number of causal genes fixed. Similarly, and as shown in the bottom left panel, the performance of non-linear `networkGWAS` is robust with respect to tuning the signal from purely linear to purely non-linear, while keeping the SNR, RCN, *r*_c_ and *n*_cg_ identical to those of scenario S0, again outperforming all comparison partners across the entire range of RLNs. Remarkably, the performance of linear `networkGWAS` starts to differ from that of its nonlinear counter-part, only for signals which are dominantly non-linear and pars the non-linear `networkGWAS` up until an RLN of 37.5: 62.5. This is in line with observations made elsewhere: approaches designed to detect statistical significance of single loci will miss those with modest marginal effects and large interactions [18]. Lastly, the strong performance of `networkGWAS` and its dominance over the comparison partners breaks down as we tune the ratio of causal neighbors from ~ 0.8 to 0.0 while keeping the SNR, RLN, *r*_c_ and *n*_cg_ the same as in scenario S0. This is shown in the bottom right panel of Figure 2, and corresponds to a gradual transition from a few, large causal subgraphs in the PPI network, via multiple medium-sized causal subgraphs, to many isolated causal genes. In the latter case, the signal–by construction–is independent of the PPI network structure, and hence that structure cannot be exploited by any network-guided method to enhance performance. The reduction of the ratio of causal SNPs on a causal gene does not decrease the performance of any method since the SNR remains constant, as visible in the Figure S.1, left panel. Instead, when the signal is concentrated on a fewer number of genes, i.e., when we reduce the number of causal genes while maintaining the SNR unvaried, the performance of `networkGWAS` remains untouched, while FaST-LMM-Set and NAGA drastically improve, suggesting that incorporating the biological network structure can substantially improve the results when the signal is spread across a high number of causal genes (e.g., 50), as it is for complex traits. These results are reported in Figure S.1.

## 4 Applications

To illustrate `networkGWAS`’s ability to allow for the discovery of new statistically significant genotype-phenotype associations, we apply `networkGWAS` to natural phenotypes from *A. thaliana* and *S. cerevisiae*. Specifically, we study these phenotypes with `networkGWAS` and its comparison partners, i.e., NAGA, dmGWAS, FaST-LMM-Set, and we additionally employ traditional univariate GWAS [15]. To identify statistical associations in presence of *p*-values, we use the hierarchical procedure [21] detailed in Section S.5, which allows to correct for multiple testing while controlling the FDR, whose level we set at 0.05. In the absence of a ground truth, we evaluate the findings of our method by (i) comparing our results to already published studies, (ii) comparing the gene neighborhoods identified as significantly associated by our method with the genes identified by the comparison partners, and (iii) investigating the potential biological relevance of identified genes in processes related to the phenotype.

### 4.1 Application to *A. thaliana*

When searching for associations with respect to natural phenotypes, we continue to use the *A. thaliana* genotype data, but rely on the larger STRING database for the PPI network [26]. The phenotypes are selected from the AraPheno [24] database (see Section S.6.2 for details).

For most of the phenotypes, there are no statistically significant associations found by any method. However, for phenotype 704, the univariate GWAS alone returns statistically significant SNPs, which are located on seven genes not connected through the PPI network (see Section S.9.1 for more detail). This scenario where the association signal is located on genes not connected through the PPI network is reminiscent of the simulation setting S6, where `networkGWAS` suffers since it cannot exploit the biological network information. `networkGWAS`, as already mentioned, does not report statistically significant associations for the analyses performed on the selected *A. thaliana* phenotypes. While this results in having no new biological insights underlying these studied complex traits, having a statistically sound approach to obtain a *p*-value that allows one to define which neighborhoods/subnetwork to consider as associated with the phenotype is an actual advantage compared to the considered network-based baselines. In contrast, methods that solely provide a ranking have no means to indicate to the practitioner that there is no significant association. The studied natural phenotypes might simply not present a genetic basis or, if they did, it might be enclosed in subnetworks not included in the current analysis, either for the lack of current knowledge of the employed biological network or for the choice of the type of biological network itself. Note that to date there are no significantly associated genetic variants reported on the AraGWAS Catalogue for the analyzed phenotypes.

### 4.2 Application to *S. cerevisiae*

In addition to *A. thaliana*, we test our method on *S. cerevisiae*. Both the GWAS dataset and the phenotypes have been obtained from the study by Peter *et al*. [20]. The PPI network has been downloaded from the STRING database [26]. More detail in Section S.6.3.

Polynomial `networkGWAS` reports no statistically significant findings, whilst linear `networkGWAS` returns two statistically significant neighborhoods on the phenotype YPGALACTOSE, which represents the growth ratio between the stress condition where the growing medium contains galactose instead of glucose and the standard growing condition for *S. cerevisiae* (see Table S.3 for further details). The two neighborhoods identified by linear `networkGWAS`, namely *YGR252W*’s and *YNL031C*’s, have 74 overlapping genes, and their union is composed of 265 genes in total (listed in Section S.9.2), representing a unique connected subgraph. We analyze these findings from different perspectives. As a preliminary investigation, we contrast them with what has been surfaced by the comparison partners and with the results presented in the original study [20]. As for p-value providing methods, FaST-LMM-Set (i.e., the analysis of genes) reports no statistically significant findings for any of the phenotypes, while traditional univariate GWAS is able to identify statistically associated SNPs for nine of them (see Table S.4 for a comparison with what presented by Peter *et al*.). For YPGALACTOSE, however, the latter returns no associations. Since NAGA and dmGWAS do not provide *p*-values, but rather scores, we identify the associated genes and subnetworks for YPGALACTOSE by following the strategies proposed by the respective authors, namely selecting the first 100 top-ranked genes for NAGA and the top 1% subnetworks for dmGWAS. This leads us to have 100 genes for NAGA and 74 genes for dmGWAS, among which 10 (7 from NAGA and 5 from dmGWAS, with two overlapping gene) are in common with `networkGWAS`’ findings. When comparing linear `networkGWAS`’ findings with the traditional GWAS reported in [20], we found no overlap between the 265 genes surfaced by `networkGWAS` and the 5 genes reported by this study. The latter are isolated genes according to the PPI network, hence violating the fundamental assumption of network-based methods designed for finding interacting genes that collectively carry association signal.

To biologically interpret `networkGWAS`’ obtained results on YPGALACTOSE, we rely on the PANTHER Classification System [27]. Specifically, we use the PANTHER Over-representation Test (PANTHER version 17.0), with PANTHER GO-slim Biological Process as the annotation dataset and Fisher’s exact test with FDR correction as the test type. We find that the genes in the 265-gene subgraph identified by `networkGWAS` are significantly enriched (the maximum *p*-value is 4.70e-02) in processes related to (i) DNA replication, (ii) chromatin organization, (iii) the cell cycle, and (iv) DNA transcription. All of these categories of processes are known to be affected when the *S. cerevisiae* organism undergoes a stress condition [7, 19]. This demonstrates that `networkGWAS` is capable of identifying neighborhoods of genes that are involved in biological processes related to the analyzed phenotype. We perform the same over-representation analysis on the associations found in the original study [20] on YPGALACTOSE; there are, however, no statistically enriched processes.

We furthermore explore the identified associated neighborhoods by means of a *post hoc* linear association. We begin with an univariate association analysis on the 3,549 SNPs included in the two neighborhoods, which led to no significance. Motivated by this result, we hypothesized the signal being of multivariate nature, and performed a Lasso analysis. We obtained the signal coming from 32 SNPs located on 25 genes (Table S.5). Interestingly, the locations of these genes are distributed across 10 different chromosomes, which highlights the benefits of including the PPI network information as a means to lead the creation of the sets of SNPs to test.

## 5 Discussion

We have defined a principled way to perform gene based genome-wide association studies utilizing network information as a prior in the process of testing statistical associations, allowing us to directly and in a statistically rigorous manner obtain *p*-values for entire PPI-based gene neighborhoods, which represent biological pathways. This conceptually differs from the state-of-the-art network-based GWAS methods, which use the PPI information as a way of post-selection, hence resulting in absence of statistically sound *p*-values. We have demonstrated the superior performance of our PPI-network based SNP set-based test, `networkGWAS`, compared to state-of-the-art SNP-set based methods [17] and approaches that incorporate PPIs [6, 29]. Moreover, we have done so in a wide range of simulation settings for rare and common variants including very low SNRs, i.e., very low heritability, different numbers of SNPs/genes carrying the association signal, and various mixtures of linear and non-linear signal. Concerning the latter, it is worth noting that none of the comparison partners can incorporate an explicit search for SNP interactions significantly associated with the phenotype, which our method–by means of employing a non-linear SNP-set kernel–is capable of. `networkGWAS` is only outperformed if our underlying assumption, that the SNPs in neighborhoods of interacting genes are collectively related to the phenotype of interest, is strongly violated.

We furthermore have employed `networkGWAS` to study various phenotypes of *S. cerevisiae* and *A. thaliana*. On the *S. cerevisiae* phenotypes, `networkGWAS` finds collectively significantly associated genes that were almost entirely undiscovered by its strongest competitor, NAGA, demonstrating the complementarity of the methods. In addition, it is particularly noteworthy that `networkGWAS` is capable of finding biologically plausible associations when the single-gene based SNP-set method does not, highlighting the value of incorporating the PPI network information. When analyzing the *A. thaliana* phenotypes, instead, `networkGWAS` finds no statistically associated neighborhoods. Although this might be perceived as a discouraging result, the studied phenotypes do not necessary present a genetic basis or association signal in the selected searching space. Hence, the fact that `networkGWAS` provides a statistically sound approach to identify the presence–if any–of significant associations is an actual advantage in comparison to the network-based comparison partners, which would output the top genes or subnetworks even on a phenotype presenting noise rather than actual signal. Furthermore, the presence of *p*-values allows to rigorously account for multiple testing correction.

As already pointed out, the application of `networkGWAS` on *S. cerevisiae* exemplifies the benefit of exploiting the biological network information. However, PPIs supported by multiple evidence, i.e., high confidence interactions, are not available (yet) for all the genes of a particular organism. For example, for *A. thaliana* only ~ 56% of the genes participate in known high-confidence PPIs, while for *S. cerevisiae*, high confidence interactions are known for ~ 92% of the genes. This may explain the results on *A. thaliana* in the sense that the current PPI network is likely incomplete and potentially associated neighborhoods of genes have not yet been identified as forming a neighborhood due to yet undiscovered interactions. Fortunately, as the knowledge of biological pathways and gene-gene interaction increases, more and more evidence will be gained and PPIs discovered, thus further approaching the state of complete PPI networks. Another aspect to consider is that `networkGWAS` can only discover the collective signal of SNPs which can be mapped onto genes. This drawback can be addressed by additionally applying traditional methods, such as univariate GWAS, to such SNPs, benefiting from increased test power due to a reduced search space.

`networkGWAS` provides *p*-values for the association of entire gene neighborhoods, and cannot single-out precise genes or SNPs within such neighborhoods as more or less strongly contributing to that association signal. If one is interested in exploring this aspect, *post hoc* analysis needs to be performed and should be guided by the kernel *K*_s_ chosen in `networkGWAS`. I.e., when a linear kernel was applied, one linear *post hoc* analysis is appropriate, as is done for the findings obtained on YPGALACTOSE. When, instead, a polynomial kernel was applied, the associated neighborhoods present nonlinear signal and epistasis search represents a viable route for *post hoc* analysis.

A potential limitation of the approach presented here lies within the choice of testing one-hop neighborhoods. While testing only direct interactions of a given gene is meaningful from a biological perspective, we acknowledge that the neighborhood depth technically constitutes a choice of hyperparameter. While the use of *k*-hop neighborhoods for *k* ≥ 2 needs to be investigated in the future, based on our simulations we are confident that at least in medium-to-high RCN scenarios, `networkGWAS` is already capable of detecting signal that is spread further than across the 1-hop neighbors of a causal center-gene. This is supported by `networkGWAS`’ findings on the *S. cerevisiae* phenotype YPGALACTOSE, where the two statistically associated neighborhoods are connected through the PPI network. We note that even if *k* ≥ 2, our method still scales linearly with the number of nodes in the network.

Another area for further research is to exploit the kernelized nature of `networkGWAS` and use more complex kernels matrices *K*_s_ in our LMM given in Equation (2). When designing such kernels, one may either (i) continue to focus on the SNP content of genes, and experiment with the type of nonlinearity, or (ii) depart from solely using SNPs as features and instead leverage the information in gene properties such as the number of minor alleles on the SNPs belonging to a gene, in combination with graph kernels or GCNs. Lastly, `networkGWAS` as presented here can be considered an instance of a more fundamental framework, which can be naturally extended and be used, for example, to study the same phenotype under different perspectives: (i) by utilizing various biological networks (e.g., gene co-expression networks), and (ii) by employing diverse ways to map the SNPs to the genes (e.g., chromatin mapping). In this sense, `networkGWAS` represents a very versatile tool.

