## Supplementary text for "networkGWAS: A network-based approach to discover genetic associations"

### Supplementary Material

#### S.1 Covariates as fixed effects

Equation (2) introduces the LMM model that `networkGWAS` relies on, and, in particular, the matrix of the fixed effects $X_{\mathrm{f}}$. The latter comprises the vector of 1s corresponding to the intercept, but it could also include other additional covariates when needed. In the context of GWAS, these covariates are factors–different from the studied genetic variants–that are expected to have an impact on the studied phenotype and, hence, that might cause spurious associations, resulting in not calibrated $p$-values. For example, when studying height or certain diseases in *H. sapiens*, it might be reasonable to include age and sex as covariates since they might affect these phenotypes. Another aspect that could influence the phenotype and cause miscalibration in the distribution of the $p$-values is the so-called population structure, which we already introduced in the main manuscript. To mitigate the effect of population structure, one could include as covariates in $X_{\mathrm{f}}$ the principal components calculated as in Price *et al.* [S20].

#### S.2 Computational cost

We base our method on the FaST-LMM-Set procedure [16], and in particular, we use the one-variance component model. A single test on a neighborhood using this procedure presents a time complexity of $O(nn_{\mathrm{s}}^2)$ in two cases, namely (i) when $K_{\mathrm{s}}$ is the linear kernel and the number of SNPs in the test set $n_{\mathrm{s}}$ is less than the number of individuals $n$, and (ii) when $K_{\mathrm{s}}$ is the polynomial kernel and $n_{\mathrm{s}}(n_{\mathrm{s}}-1)/2 < n$, because the trivial factorization in the reproducing kernel Hilbert space of dimension $n_{\mathrm{s}}(n_{\mathrm{s}}-1)/2$ has to be obtained. Otherwise, each test costs $\mathcal{O}(n^3)$. Hence, in the best-case scenarios, testing $n_{\mathrm{g}}$ neighborhoods has a runtime of $\mathcal{O}(n_{\mathrm{g}}nn_{\mathrm{s}}^2)$, where $n_{\mathrm{g}}$ is the number of genes considered in the analysis. Before applying FaST-LMM-Set, however, `networkGWAS` requires the aggregation of gene neighborhoods which scales linearly in the number of edges of the PPI employed. As biological networks can be assumed to be only sparsely connected, this leads to a runtime of order $\mathcal{O}(n_{\mathrm{g}})$. Secondly, in order to obtain $p$-values from the test statistics, we have to repeat both the neighborhood aggregation and the $n_{\mathrm{g}}$ set tests for each of the $n_{\mathrm{p}}$ permutations. This results in a computational cost of $\mathcal{O}\left(n_{\mathrm{p}}n_{\mathrm{g}}nn_{\mathrm{s}}^2\right)$ if $n_{\mathrm{s}} < n$ for the linear kernel or $n_{\mathrm{s}}(n_{\mathrm{s}}-1)/2 < n$ for the quadratic kernel. Otherwise, the cost amounts to $\mathcal{O}\left(n_{\mathrm{p}}n_{\mathrm{g}}n^3\right)$. Therefore, while the cubic scaling is not favorable for large cohort sizes, asymptotically, run time scales linearly in $n$. For the *S. cerevisiae* data, the median number $n_{\mathrm{s}}$ of SNPs per neighborhood is 215. Hence, for the majority of *S. cerevisiae* PPI neighborhoods tested, the cubic-to-linear speed up sets in for cohort sizes as small as $n = 971$ for linear `networkGWAS`, while for non-linear `networkGWAS` that speed-up would be achieved for cohorts of $n \gtrsim 471,000$ samples. Note that considering the computational time to perform the permutations themselves, i.e. rotating the SNPs and permuting the network, would add $O(n_{\mathrm{p}})$ and $O(n_{\mathrm{p}}n_{\mathrm{g}})$–which becomes $O(n_{\mathrm{p}}n_{\mathrm{g}}^2)$ in case of non-sparse network–to the total time complexity respectively, terms that are dominated by the above derived time complexity, regardless of the case.

#### S.3 Model choices: motivation

The log-likelihood of the model underlying the original FaST-LMM-Set [S14,S15] method writes:

$$LL(\beta, \sigma_{\mathrm{e}}^2, \tau) = \log \mathcal{N}(\vec{y}|X\vec{\beta}; \sigma_{\mathrm{e}}^2 I + \sigma_{\mathrm{g}}^2[(1-\tau)K_{\mathrm{c}} + \tau K_{\mathrm{s}}])\,, \tag{S.1}$$

and features two random effects: one to capture confounders ($K_{\mathrm{c}}$) and another to account for similarity among the SNPs of the set to be tested ($K_{\mathrm{s}}$). The covariance matrices are defined as $K_{\mathrm{c}} = \frac{1}{n_{\mathrm{c}}}V_{\mathrm{c}}V_{\mathrm{c}}^T$ and $K_{\mathrm{s}} = \frac{1}{n_{\mathrm{s}}}V_{\mathrm{s}}V_{\mathrm{s}}^T$, where $V_{\mathrm{c}}$ contains the $n_{\mathrm{c}}$ SNPs from which relatedness is estimated and $V_{\mathrm{s}}$ contains the $n_{\mathrm{s}}$ SNPs to test. As introduced in [17], the parameter $\tau \in [0,1]$ serves to distinguish the null model (i.e., $\tau = 0$) from alternative models (i.e., $0 \leq \tau \leq 1$), and is estimated from the GWAS data set by means of restricted maximum likelihood. Once the value of equation (S.1) has been calculated for the null and the alternative hypotheses, it is possible to compute the likelihood-ratio test statistics, i.e., $t = -2(LL(\tau = 0) - LL(0 \leq$

$\tau \leq 1$)). Now, the original FaST-LMM-Set follows the spirit of Wilks' theorem [S31] and results by Greven *et al.* [S9] and employs a mixture of $\chi^2$ distributions,

$$p_0(x) = a\chi_0(x) + b\chi_d(x) \tag{S.2}$$

to serve as parametric distribution of the test statistic under the null hypothesis. The parameters are determined by fitting $p_0(x)$ to the 10% most significant tail of the null distribution of test statistics obtained by permuting individuals for only the SNPs in the set of interest, that is, permuting the rows of $V_{\text{s}}$.

We chose to implement an alternative permutation strategy for the following reasons. As already mentioned above, $K_{\text{c}}$ is the kernel matrix that captures confounders, and represents the genetic similarity matrix (GSM), which measures and corrects for population structure in the form of a realized relationship matrix (RRM) [S8,S10]. $K_{\text{c}}$ is calculated as $\frac{1}{n_{\text{c}}}V_{\text{c}}V_{\text{c}}^T$; the standard, computationally efficient way to choose the SNPs to include in $V_{\text{c}}$ is the so-called "leave-one-out-chromosome" strategy [S13], where the sets of SNPs on each chromosome are tested separately and $K_{\text{c}}$ is constructed using the SNPs located on the other chromosomes. Such a strategy is not computationally reasonable when testing neighborhoods, as genes interacting through a biological network can be, and frequently are, located on different chromosomes. Therefore, when excluding the GSM $K_{\text{c}}$ for such reasons from our model, employing FaST-LMM-Set permutation scheme, i.e., permuting the individual of $K_{\text{s}}$, would result in not preserving the population structure signal when generating the null distribution [S11]. Hence, we implemented the permutation scheme described in Section 2.3, which allowed us to preserve population structure when constructing the null distribution when testing neighborhoods.

Furthermore, while fitting a parametric null distribution saves computation time by decreasing the number of permutations needed, we found the resulting estimate of the distribution $p_0(x)$ of test statistics under the null hypothesis did not reflect the true distribution in our case. Similar limitations of the above parametric approach in LMMs have been discussed in [S19], and hence we adhere to a non-parametric distribution for `networkGWAS` instead, enabling us to obtain well calibrated $p$-values across scenarios and simulations, as exemplified in Fig. S.3. More specifically, `networkGWAS` allows us to obtain a genomic inflation factor $\lambda_{\text{GC}} \in [0.8, 1.2]$ for 98% of the scenarios analysed in the *A. thaliana* semi-simulation, in 92% of the natural *A. thaliana* phenotypes, and in 100% of the *S. cerevisiae* phenotypes studied. As anticipated in Section 2.3, for some of the simulated phenotypes in the *H. sapiens* use case, the portion of high $p$-values present deflation, leading to $\lambda_{\text{GC}} \notin [0.8, 1.2]$, while the rest of the distribution of the $p$-values, including the low-$p$-values range, look calibrated. Note that we obtained these results without any hyperparameter selection procedure. In fact, our permutation technique requires only one parameter to set, the percentage of edges to permute, which we set to 50%. In contrast, the original FaST-LMM-Set requires an optimization step aimed at selecting the SNPs modeling the relatedness in the dataset, i.e., the SNPs to include in $V_{\text{c}}$. In fact, to yield the computational savings that reside at the core of the FaST-LMM procedure [S13], the sum of the dimensions of the kernel matrices $K_{\text{c}}$ and $K_{\text{s}}$ must be kept well below the number $n$ of samples studied. To this end, the GSM $K_{\text{c}}$ is constructed from a limited number of SNPs chosen on the basis of their $p$-values in an uncorrected linear regression with respect to the phenotype of interest, following the leave-one-out-chromosome approach. Lastly, to identify the precise number of SNPs to be included in the GSM, the latter is constructed with an increasing number of SNPs–starting with those associated with the lowest $p$-values–until the first minimum in the genomic inflation factor $\lambda$–defined as the ratio of the median observed to median theoretical test statistic–is met [S13].

#### S.4 Number of permutations in `networkGWAS`

As detailed in Section 2.3, we obtain our null distribution $T_0$ by pooling the statistics from each neighborhood from each permutation. Hence, if $n_{\text{g}}$ is the number of neighborhoods and $n_{\text{p}}$ is the number of permutations, the non-parametric null distribution is composed of $n_{\text{g}} \times n_{\text{p}}$ tests statistics, decreasing the total number of permutation required. A question now arises, namely "how to define a proper number of permutations $n_{\text{p}}$?". There are two criteria that allow one to answer this question. First, one could check the distribution of the null test statistics obtained after each permutation and find after which permutation this distribution stabilizes. Second, our method foresees this hierarchical procedure based on B-H as a means to correct for multiple testing. Hence, the minimum obtainable $p$-value, which depends on the number of permutations, should be “small enough” to allow for discovering significance (if any). In practise, we found that a number of permutations among 100 and 300 is a good compromise to reduce the required computation and have a valuable estimate of the $p$-values.

#### S.5 Hierarchical testing procedure

Correction for multiple testing is necessary when dealing with a multitude of hypotheses, such as in GWAS, where millions of genetic variants are studied and tested at once. In such scenarios, the false discovery rate (FDR) is often used as a measure of global error. FDR is defined as the expected proportion of findings, i.e., rejected null hypotheses, for which the null hypothesis is actually true. As defined in Section 2.3, the null hypothesis $H_j$ for `networkGWAS` signifies that the $j$-th neighborhood $\mathcal{N}_k(v_j)$ does not affect the phenotype. When studying $P$ related phenotypes (for example phenotypes obtained from the same study), we can rewrite the null hypothesis as $H_{jt}$, and it represents that the $j$-th neighborhood does not affect the $t$-th phenotype. This results in having a collection of null hypotheses $\{H_{jt}, j = 1, \ldots, M; t = 1, \ldots, P\}$. Having all these hypotheses from different neighborhoods and phenotypes, one possibility would be to control the global FDR, e.g., applying the Benjamini-Hochberg (B-H) procedure [S2] on the pooling of the hypotheses. However, this approach has been shown to present a suboptimal behaviour, that is, for phenotypes presenting no or low association signal, then the FDR would not be controlled at the defined level $q$, while for phenotypes showing medium or high association signal, this procedure would cause a decrease in the power [S6]. Peterson *et al.* [S18] define a rigorous procedure aimed at correcting the aforementioned situation, which will be detailed in the following. Consider again the collection of null hypotheses $\{H_{jt}, j = 1, \ldots, M; t = 1, \ldots, P\}$. Among them, we can identify subgroups of hypotheses, based on what we are interested in studying. In our case, one group corresponds to all the hypotheses that relate to one phenotype, and we define each group as $\mathcal{P}_t = \{H_{jt}, j = 1, \ldots, M\}$. Note that when introducing the concept of group/subgroups in the null hypotheses collection, Peterson *et al.* [S18] refer to them as “families”; here we use these terms interchangeably. From these families, we can obtain a global null hypothesis, which in our case writes $H_{\bullet t} = \cap_{t=1}^{M} H_{jt}$ and means that no neighborhood affects the $t$-th phenotype. Having defined the notation, we can introduce the procedure, which is comprised of three steps, namely:

1. Employing Simes’s method [S25] to calculate a $p$-value per each global hypothesis, i.e., $H_{\bullet t}$. Precisely, these $P$ $p$-values ($p_{\bullet t}$) are calculated as:

$$p_{\bullet t} = \min_{j=1,\ldots,M} M p_t(j)\,, \tag{S.3}$$

   where $p_t(j)$ is the $p$-value obtained for the neighborhood $j$ for the phenotype $t$.
2. Applying the B-H procedure, with FDR target set to $q_1$, on the $p$-values obtained according to point 1. The set of phenotypes for which the null hypothesis $H_{\bullet t}$ has been rejected is defined as $\mathcal{S}$.
3. For the phenotypes in $\mathcal{S}$, applying B-H on their respective subgroup of hypotheses, $\mathcal{P}_t$, with FDR target set to $q_1 * |\mathcal{S}|/P$. This step returns the statistically associated neighborhoods for the phenotypes in $\mathcal{S}$, i.e., those phenotypes that reject the null hypothesis of no neighborhood affecting them.

When Simes’s $p$-values are valid [S22], the described procedure guarantees the control of the FDR for the discovery of the phenotypes (vFDR in [S18]), and when B-H is applied on the set of global hypotheses, then it controls FDR at level $q_1$, allowing to identify $\mathcal{S}$ while indeed controlling the FDR. Then, it controls the the average FDR on the families in $\mathcal{S}$ (sFDR in [S18]), at level $q_2$, by applying B-H as described in point 2 above.

#### S.6 Data

##### S.6.1 *A. thaliana*: semi-simulation

We use the genotype dataset available on the manually curated and standardized GWAS catalog for *A. thaliana*, i.e., AraGWAS [S30], and in particular, we utilize the fully-imputed dataset which comprises 2,029 samples with 10,709,466 SNPs each. We filter the SNPs for at least 5% minor-allele frequency, leaving us with 1,763,004 SNPs. Out of those, 37,458 can be mapped to the 1,327 genes included in the PPI network, obtained from The Arabidopsis Information Resource (TAIR) database [S12] using the strict (i.e., without windows of base pairs) positional mapping of the genes downloaded from TAIR. We chose the TAIR network for our semi-simulated setting since its smaller size allowed us to run a large number of fast experiments.

##### S.6.2 *A. thaliana*

As reported in the main text, when studying natural phenotypes from *A. thaliana*, we again use the genotype data from the AraGWAS Catalog [S30], but rely on the larger STRING database for the PPI network [S26], considering the high confidence PPIs. In fact, the interactions in the STRING PPI network are annotated with one or more 'scores' which are indicators of confidence, i.e. how likely STRING judges an interaction to be true given the available evidence. These scores rank from 0 to 1, with 1 being the highest possible confidence. For our analysis we only consider high-confidence interactions with a score larger than or equal to 0.7. The phenotypes are selected from the AraPheno [S23] database, a central repository of population-scale phenotypes for *A. thaliana* inbred lines, based on the criteria that they are quasi-continuous (such as stem length, and flower diameter) and be known for at least 500 out of the 2,029 *A. thaliana* samples that we use the genotypes of. This leads to the selection of 13 phenotypes, summarized in Table S.2. For each phenotype, we collect the fully-imputed genotypes provided by AraGWAS for which the phenotype of interest has been determined. We then filter for at least 5% minor-allele frequency among those samples which–depending on the phenotype–leaves us with 1,837,440 $\pm$ 27,477 remaining SNPs. Out of those, 968,620 $\pm$ 16,200 could be positionally mapped [S12] to the 31,568 $\pm$ 18 *A. thaliana* genes that present at least one SNP mapped to it. Note that isolated genes according to the STRING PPI network are considered in the analysis.

##### S.6.3 *S. cerevisiae*

When studying *S. cerevisiae*, we obtain both the GWAS dataset and the phenotypes from the study by Peter *et al.* [S17]. The GWAS dataset comprises 1,011 *S. cerevisiae* isolates of 83,794 genetic variants (82,869 SNPs and 925 copy-number variants, MAF $\geq$ 0.05) each. For 971 of the 1,011 strains, the growth ratio between 35 stress conditions and the standard growing condition for *S. cerevisiae* has been recorded. Among these 35 phenotypes, we consider the 23 that present a Gaussian distribution of the values, listed in Table S.3. The *S. cerevisiae* PPI network has been downloaded from the STRING database [S26], and we again only consider high confidence interactions, i.e., interactions with confidence score greater or equal than 0.7, in order to limit the bias due to the noise/uncertainty of the PPIs. After performing the SNPs-genes strict positional mapping [S3], 59,787 SNPs and 5,737 genes remain.

#### S.7 Phenotypes simulation

As mentioned in the main text, to simulate the phenotypes, we firstly define the following parameters: (a) the number of genes $n_{\mathrm{cg}}$ that carry causal SNPs, henceforth called causal genes, (b) the ratio of causal SNPs on a causal gene, $r_{\mathrm{c}}$, (c) the mean ratio of causal neighbors (RCN) of a causal gene, (d) the signal-to-noise-ratio (SNR), and (e) the mixing ratio of linear-to-nonlinear (RLN) signal. All the above defined parameters are systematically varied in our experiments. Thus, we investigate the robustness of our method's and comparison partners' performance as we depart from the most amenable scenario (S0) of a purely linear signal, spread across a single or very few causal subgraph(s) composed of a high number of causal genes, with a high ratio of causal neighbors, a realistic signal-to-noise ratio, and a high ratio of causal SNPs on the causal genes. Table S.1 summarizes the scenarios simulated, starting from our anchor scenario S0 and deviating from its conditions by:

- Varying the SNR while keeping the RCN, RLN, $r_{\mathrm{c}}$, and $n_{\mathrm{cg}}$ constant,
- Varying the RLN while keeping the SNR, RCN, $r_{\mathrm{c}}$, and $n_{\mathrm{cg}}$ constant,
- Varying the RCN while keeping the SNR, RLN, $r_{\mathrm{c}}$, and $n_{\mathrm{cg}}$ constant,
- Varying the $r_{\mathrm{c}}$ while keeping the SNR, RCN, RLN, and $n_{\mathrm{cg}}$ constant,
- Varying the $n_{\mathrm{cg}}$ while keeping the SNR, RCN, RLN, and $r_{\mathrm{c}}$ constant.

We thereby study the isolated effects of moving towards more challenging/different RCN, RLN, SNR, $r_c$, and $n_{cg}$ respectively.

In order to realize the predefined scenarios, we follow the procedure below for the simulation of the artificial phenotype:

1. Choice of causal genes:
   (a) For RCN equal to 0./0.4/0.8, randomly select $n_{cg}$/5/1 node(s) in the network and define them as causal.
   (b) Randomly include 0%/40%/100% of the $k$-hop neighbors of the starting node(s), then 0%/40%/100% of the $k$-hop neighbors of the newly included nodes and so on, until the pre-defined number $n_{cg}$ of causal genes is reached.
   (c) Should the starting node(s) belong to a disconnected subgraph(s) smaller than as to allow for the inclusion of $n_{cg}$ causal genes, repeat (1)-(3) until $n_{cg}$ causal genes have been defined.
   (d) Compute RCN, and if RCN is not in $[0.]$ / $[0.4, 0.5]$ / $[0.8, 1.0]$, disregard and repeat (1)-(4).
2. Choice of causal SNPs and causal interactions:
   (a) Randomly select a ratio of $r_c$ of the SNPs of each causal gene as carrying the signal.
   (b) For each causal subgraph, define one of the nodes with highest degree as the center of the subgraph. For each causal SNP belonging to a non-center node of the same causal subgraph, randomly select a causal SNP on the center node and define the interaction between the two SNPs as causal. Ensure that each center-node causal SNP has at least one interaction partner.
3. Computation of the artificial phenotype:
   (a) Simulate the phenotype as
   $$\vec{y} = cX \cdot \vec{\beta} + (1-c)X^{(2)} \cdot \vec{\beta}^{(2)} + \vec{\epsilon} \tag{S.4}$$
   where $X$ is the $n$ by $p$ matrix of all SNPs across the genes considered in the analysis (i.e., 37,458 in the semi-simulated case and 10,000 in the simulated case), $\vec{\beta}$ represents the fixed effect of these SNPs, $\vec{\epsilon}$ models the noise, $X^{(2)}$ is the $n$ by $p(p-1)/2$ second-order design matrix of all SNP interactions, $\vec{\beta}^{(2)}$ comprises the fixed effects of all SNP interactions, and the coefficient $c$ serves to tune between the ratio of linear to non-linear signal. $\vec{\epsilon}$ is drawn from a multivariate normal distribution $\mathcal{N}(\vec{0}, I)$.
   (b) Set $\beta_i$ to a fixed and positive value $b$ if the $i^{\text{th}}$ SNP is a causal one, or to zero otherwise. Choose $b$ such that for $c = 1$, the SNR equals the desired value.
   (c) Set $\beta_i^{(2)}$ to a fixed and positive value $b^{(2)}$ if the $i^{\text{th}}$ SNP-interaction is a causal one, or to zero otherwise. Choose $b^{(2)}$ such that for $c = 0$ the SNR equals the desired value.
   (d) Choose $c$ such that the desired RLN is realized.

Tuning the signal-to-noise ratio (SNR) in our simulation allows us to mimic a range of heritability values $h$ by means of the relation SNR $= h/(1-h)$. With low but realistic heritability values ranging from $h = 0.1$ to $h = 0.4$, SNRs to be investigated stretch from SNR $= 0.1$ to SNR $= 0.67$, which includes all the values investigated by us. Lastly, note that we create 5 random realizations of each scenario S0-S10, in order to estimate the variance in performance.

#### S.8 Simulated rare variants and phenotypes in *H. sapiens*

Having assessed `networkGWAS`'s ability of successfully detecting known associations on the *A. thaliana* common variants semi-simulated use case, we decided to test `networkGWAS` on yet a different scenario, namely on synthetic rare variants and phenotypes from a different organism, i.e., *H. sapiens*.

##### S.8.1 Experimental setup

With the aim of simulating rare variants with realistic LD and MAF distributions, we use the sim1000G package [S5], which requires the variant call format (VCF) file of the genomic region of interest as the unique input. We choose the VCF file for chromosome 10 from Phase III 1000 genomes sequencing data [S1], and we use PLINK [S21,S24] to extract the variant calls for the European population. We employ bcftools [S4] to further manipulate the VCF file, namely to filter out the variants that are not mapped – according to the MyGene database [S32,S33] – onto the 398 interacting genes through the human STRING PPI network [S26]. This is always restricted to chromosome 10. Note that working on a subset of the *H. sapiens* genes does not compromise the results since the genotype-phenotype association signal is simulated. Having obtained the desired VCF file, we therefore use sim1000G to simulate 10,000 rare variants, i.e. MAF $\leq 0.1$, from 500 unrelated subjects. The generation of the synthetic phenotypes follows the same steps and parameter choices as detailed in Section 3.1, apart from $n_{cg}$, which is set to 15, 8 or 2 to have the same causal/non-causal genes ratio as for the *A. thaliana* use case. Lastly, the simulation settings S0 to S10 are chosen following the common-variant semi-simulations listed in Table S.1. Comparison partners and performance evaluation follow what reported in Section 3.2.

##### S.8.2 Results

The results obtained for the rare-variant simulations are summarized in Figure S.2 and are very similar in nature to the results obtained on common variants. As can be seen in the top left panel, in our scenario S0 of a purely linear signal, spread across a single or very few causal subgraph(s) with a high ratio of causal neighbors, and high signal-to-noise ratio, both linear and non-linear `networkGWAS` outperform all the comparison partners by achieving an AUPRC of 69.9%±17.9% and 65.6%±18.4%, respectively. As demonstrated in panel (b) and (c) of Figure S.2, this dominance is maintained while varying SNR and linear-to-nonlinear signal mixtures, and only vanishes for a fully nonlinear phenotype. Note, however, that in the rare variants case, `networkGWAS` is less robust to both increasing SNR and increasing nonlinearity of the signal than for the common variants. Furthermore, for RVs, the non-linear SNP-set kernel does not perform better than the linear one in any of the scenarios studied. As in the common-variant simulations, in the rare-variant case, too, `networkGWAS` cannot outperform its comparison partners in the case of isolated causal-genes, while `networkGWAS` does outperform comparison partners as soon as the PPI network holds phenotype-relevant information. This is shown in panel (d) of Figure S.2. Similar considerations to the *A. thaliana* use case apply to the variation of the ratio of causal SNPs on the causal genes and of the number of causal genes; results shown in the panels (e) and (f) of Figure S.2.

#### S.9 Statistically associated SNPs, genes and neighborhoods

##### S.9.1 Univariate GWAS hits on *A. thaliana* phenotype 704

The genes where the SNPs identified by traditional univariate GWAS analysis are located are *AT4G00630, AT4G00730, AT4G00740, AT4G00752, AT4G01010, AT5G45750,* and *AT5G45760.* We compare these findings with the results available on the AraGWAS Catalog [S30] for phenotype 704, namely the number of rosette leaves. To ensure a fair comparison, we filter the published associations using the same MAF threshold we employed, we consider only genic SNPs, and we use FDR for multiple testing correction. This leads to have no significant associations on the AraGWAS Catalogue for phenotype 704. To biologically investigate these findings, we use the gene ontology (GO) Term Enrichment for Plants [S27] provided by TAIR, which relies on the PANTHER Classification System [S16,S29]. We use the PANTHER Over-representation Test with the same settings as done for *S. cerevisiae*. This analysis does not yield to any significantly enriched process. However, the TAIR database reports that *AT4G00630*, *AT4G00740*, and *AT5G45750* are expressed in the rosette leaves, therefore representing biologically plausible results.

##### S.9.2 `networkGWAS` ($K_{lin}$) on *S. cerevisiae* phenotype YPGALACTOSE

*YAL013W, YAL019W, YAR003W, YAR007C, YBL002W, YBL003C, YBL008W, YBL023C, YBL052C, YBL088C, YBL103C, YBR034C, YBR081C, YBR103W, YBR109C, YBR112C, YBR114W, YBR133C, YBR136W, YBR160W, YBR175W, YBR195C, YBR198C, YBR202W, YBR215W, YBR245C, YBR274W, YBR289W, YCL010C, YCL061C, YCR012W, YCR033W, YCR065W, YCR082W, YCR084C, YDL003W, YDL017W, YDL042C, YDL074C, YDL139C, YDL140C, YDR013W, YDR034C, YDR052C, YDR083W, YDR096W, YDR145W, YDR167W, YDR176W, YDR181C, YDR190C, YDR191W, YDR216W, YDR217C, YDR224C, YDR227W, YDR303C, YDR334W, YDR359C, YDR392W, YDR440W, YDR448W, YDR469W, YDR477W, YDR489W, YEL009C, YEL018W, YEL021W, YEL032W, YEL056W, YER040W, YER051W,*

*YER052C, YER095W, YER110C, YER148W, YER164W, YER169W, YFL021W, YFL024C, YFL037W, YFL039C, YFR037C, YGL008C, YGL043W, YGL058W, YGL060W, YGL066W, YGL073W, YGL112C, YGL116W, YGL150C, YGL163C, YGL194C, YGL201C, YGL207W, YGL237C, YGL244W, YGL252C, YGR002C, YGR056W, YGR063C, YGR098C, YGR116W, YGR140W, YGR179C, YGR188C, YGR192C, YGR200C, YGR212W, YGR252W, YGR270W, YGR274C, YGR275W, YHL022C, YHR013C, YHR030C, YHR056C, YHR079C, YHR090C, YHR099W, YHR119W, YHR187W, YHR205W, YHR207C, YIL084C, YIL126W, YIL144W, YIL150C, YJL047C, YJL052W, YJL072C, YJL074C, YJL081C, YJL110C, YJL115W, YJL127C, YJL153C, YJL168C, YJL176C, YJL187C, YJL194W, YJR009C, YJR066W, YJR082C, YJR119C, YJR140C, YKL005C, YKL018W, YKL028W, YKL058W, YKL126W, YKR008W, YKR048C, YKR101W, YLL002W, YLL022C, YLR015W, YLR055C, YLR085C, YLR086W, YLR103C, YLR141W, YLR182W, YLR274W, YLR318W, YLR357W, YLR384C, YLR399C, YLR418C, YLR442C, YLR449W, YML010W, YML032C, YML041C, YML063W, YML069W, YML085C, YML102W, YML124C, YML127W, YMR020W, YMR033W, YMR037C, YMR044W, YMR048W, YMR075W, YMR076C, YMR127C, YMR176W, YMR223W, YMR236W, YMR263W, YMR284W, YMR312W, YNL021W, YNL031C, YNL088W, YNL097C, YNL107W, YNL112W, YNL132W, YNL136W, YNL167C, YNL206C, YNL236W, YNL246W, YNL273W, YNL312W, YNL330C, YNR023W, YOL004W, YOL006C, YOL012C, YOL051W, YOL054W, YOL068C, YOL069W, YOL146W, YOL148C, YOR001W, YOR023C, YOR025W, YOR026W, YOR038C, YOR061W, YOR064C, YOR073W, YOR123C, YOR151C, YOR194C, YOR213C, YOR244W, YOR253W, YOR290C, YOR304W, YPL001W, YPL015C, YPL016W, YPL018W, YPL042C, YPL047W, YPL082C, YPL086C, YPL101W, YPL116W, YPL127C, YPL129W, YPL138C, YPL153C, YPL181W, YPL209C, YPL235W, YPL240C, YPL254W, YPR018W, YPR019W, YPR023C, YPR031W, YPR034W, YPR068C, YPR086W, YPR133C, YPR135W, YPR161C.*

#### S.10 Figures

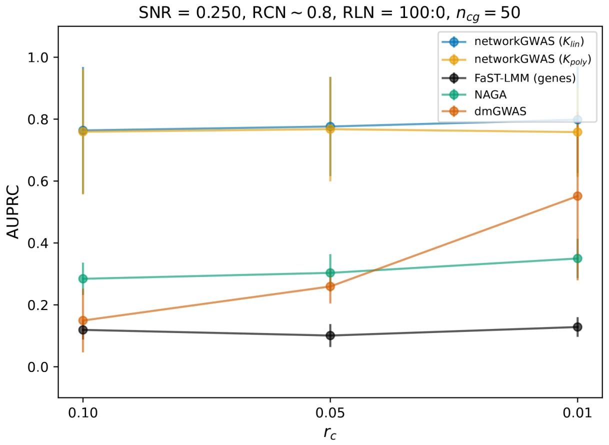


(a)

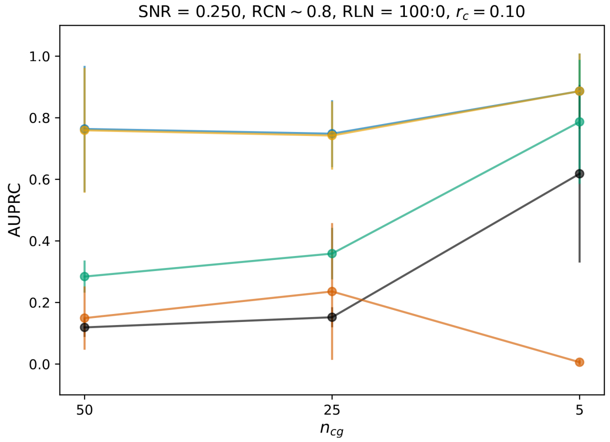


(b)

Fig. S.1: Results from simulating the phenotypes of *A. thaliana*. We present results from our method (**networkGWAS**), using either a linear ($K_{\text{lin}}$) or a polynomial ($K_{\text{poly}}$) kernel, as well as the performance of other comparison methods. We show the average AUPRC when varying when varying the ratio of causal SNPs on a causal gene ($r_{\text{c}}$) (a) or the number of causal genes ($n_{\text{cg}}$) (b), while keeping the other four variables fixed.

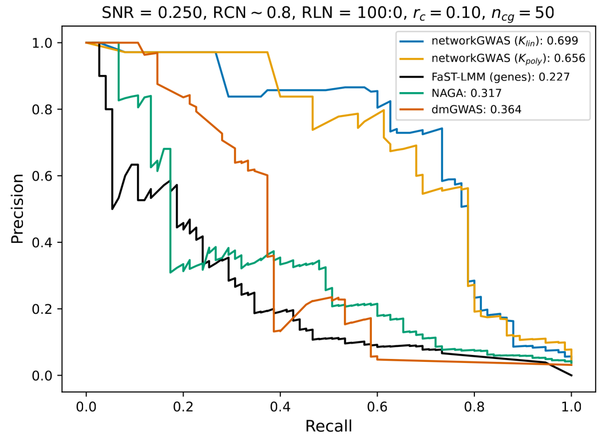


(a)

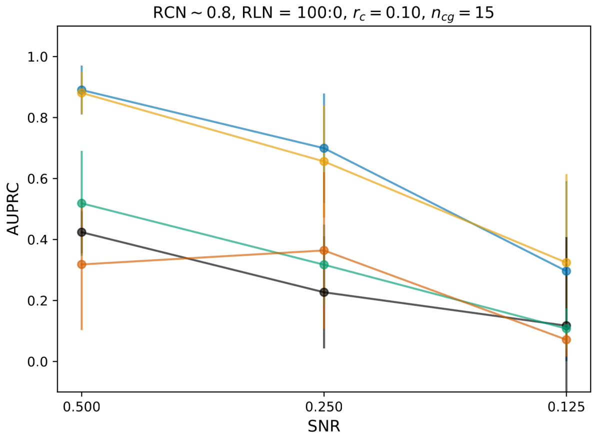


(b)

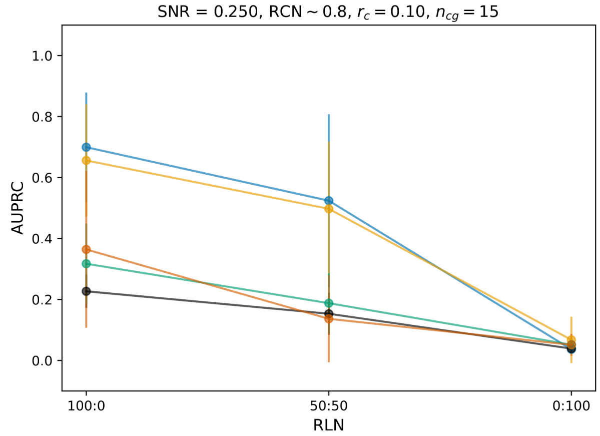


(c)

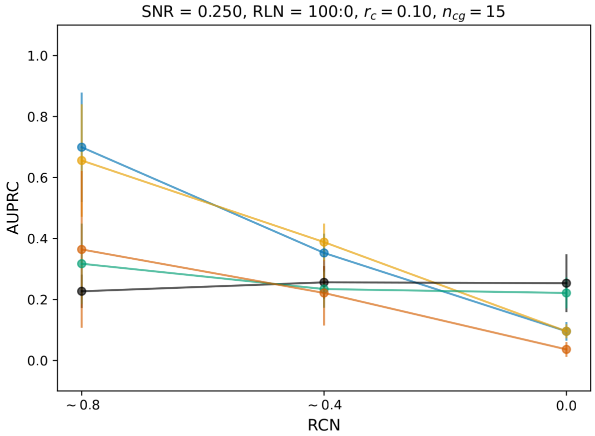


(d)

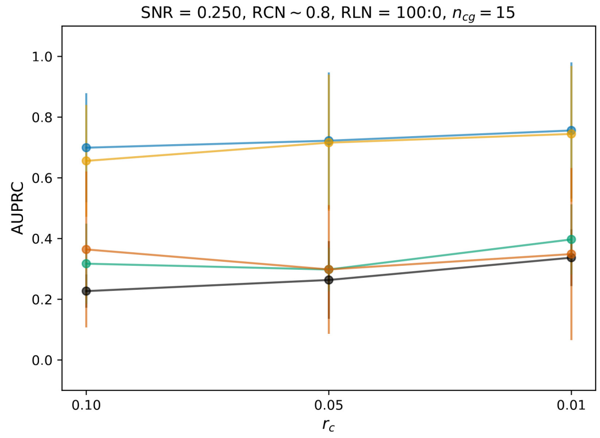


(e)

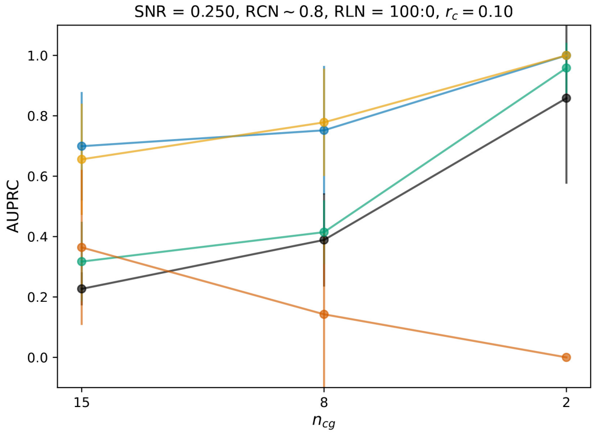


(f)

Fig. S.2: Results from our simulated *H. sapiens* use case. We present results from our method (`networkGWAS`), using either a linear ($K_{\text{lin}}$) or a polynomial ($K_{\text{poly}}$) kernel, as well as the performance of other comparison methods. In Subfigure S2a, we show the AUPRC of the baseline scenario (S0). In Subfigures 3-b-3d, we vary one variable while keeping the other four fixed: the signal-to-noise ratio (SNR) (S2b), the mixing ratio of linear-to-nonlinear (RLN) signal (S2c), the mean ratio of causal neighbors (RCN) (S2d), the ratio of causal SNPs on a causal gene ($r_{\text{c}}$) (S2e), and the number of causal genes ($n_{\text{cg}}$) (S2f).

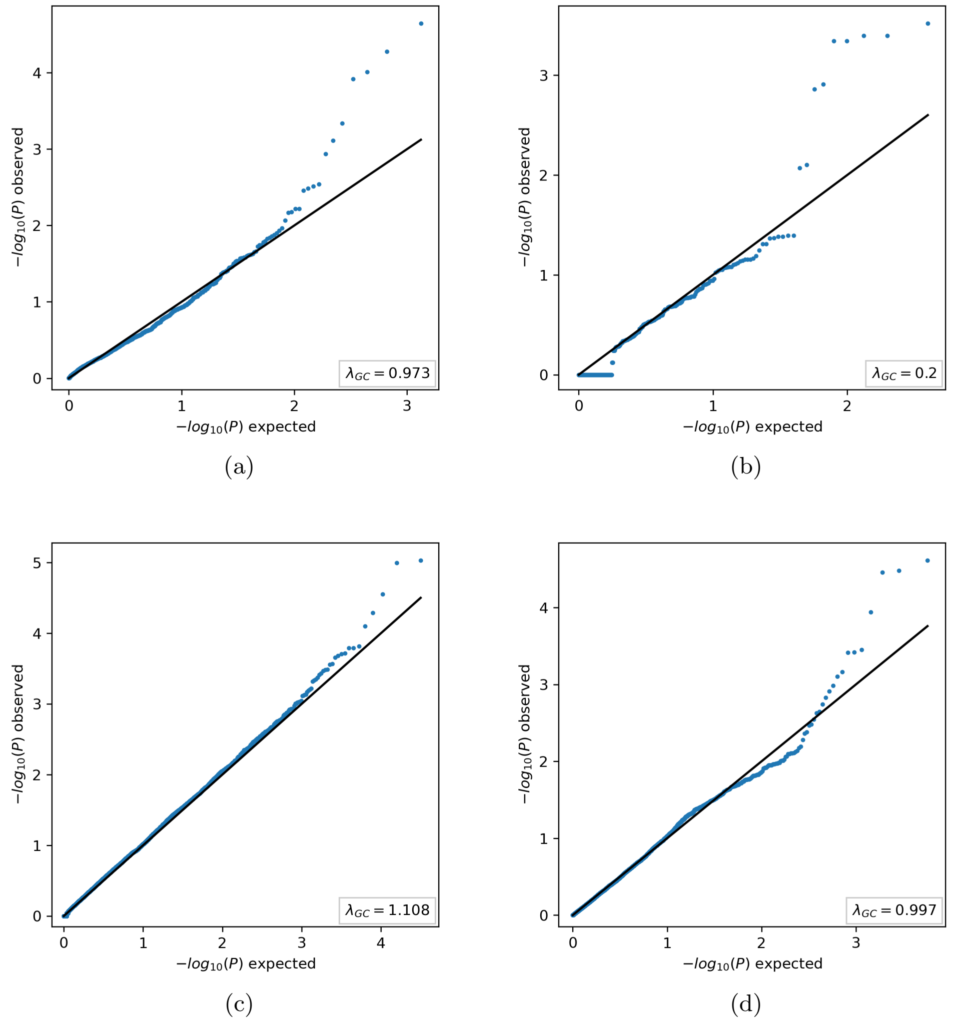


Fig. S.3: Distributions of the $p$-values obtained using `networkGWAS` on different use cases: (a) a scenario from the *A. thaliana* semi-simulations, setting S5; (b) a scenario from setting S0 for *H. sapiens* synthetic simulations; (c) the *A. thaliana* phenotype 524; (d) the *S. cerevisiae* phenotype YPGALACTOSE.

#### S.11 Tables

| | S0 | S1 | S2 | S3 | S4 | S5 | S6 | S7 | S8 | S9 | S10 |
|---|---|---|---|---|---|---|---|---|---|---|---|
| **SNR** | 0.250 | 0.125 | 0.500 | 0.250 | 0.250 | 0.250 | 0.250 | 0.250 | 0.250 | 0.250 | 0.250 |
| **RLN** | 100 : 0 | 100 : 0 | 100 : 0 | 50 : 50 | 0 : 100 | 100 : 0 | 100 : 0 | 100 : 0 | 100 : 0 | 100 : 0 | 100 : 0 |
| **RCN** | 0.8 | 0.8 | 0.8 | 0.8 | 0.8 | 0.4 | 0.0 | 0.8 | 0.8 | 0.8 | 0.8 |
| $r_{\mathbf{c}}$ | 0.10 | 0.10 | 0.10 | 0.10 | 0.10 | 0.10 | 0.10 | 0.05 | 0.01 | 0.10 | 0.10 |
| $n_{\mathbf{cg}}$ | 50 | 50 | 50 | 50 | 50 | 50 | 50 | 50 | 50 | 25 | 5 |

Table S.1: Overview of simulation settings for artificial phenotypes. All the parameters, i.e., the signal-to-noise ratio (SNR), the mixing ratio of linear to non-linear signal (RLN), the ratio of causal neighbors (RCN), the ratio of causal SNPs ($r_{\mathrm{c}}$), and the number of causal genes ($n_{\mathrm{cg}}$) are systematically varied.

| | id | phenotype | cohort size $n$ | original study |
|---|---|---|---|---|
| **N01** | 518 | relative lifetime fitness in Spain for high precipitation treatment and low density | 512 | [S7] |
| **N02** | 519 | relative lifetime fitness in Spain for high precipitation treatment and high density | 512 | [S7] |
| **N03** | 522 | relative lifetime fitness in Germany for high precipitation treatment and low density | 512 | [S7] |
| **N04** | 523 | relative lifetime fitness in Germany for high precipitation treatment and high density | 512 | [S7] |
| **N05** | 524 | relative lifetime fitness in Germany for low precipitation treatment and low density | 512 | [S7] |
| **N06** | 534 | relative seed production in Spain for high precipitation treatment and low density | 511 | [S7] |
| **N07** | 535 | relative seed production in Spain for high precipitation treatment and high density | 512 | [S7] |
| **N08** | 538 | relative seed production in Germany for high precipitation treatment and low density | 512 | [S7] |
| **N09** | 539 | relative seed production in Germany for high precipitation treatment and high density | 511 | [S7] |
| **N10** | 700 | length of main flowering stem | 680 | [S28] |
| **N11** | 704 | rosette leaf number | 850 | [S28] |
| **N12** | 705 | cauline leaf number | 904 | [S28] |
| **N13** | 707 | diameter of rosette | 656 | [S28] |

Table S.2: Overview of *A. thaliana* phenotypes.

| **id** | **media composition** | **temperature** |
| --- | --- | --- |
| YPACETATE | 2% bactopeptone; 1% yeast extract; 2% acetate; 2% agar | 30°C |
| YPD14 | 2% bactopeptone; 1% yeast extract; 2% glucose; 2% agar | 14°C |
| YPD40 | 2% bactopeptone; 1% yeast extract; 2% glucose; 2% agar | 40°C |
| YPD42 | 2% bactopeptone; 1% yeast extract; 2% glucose; 2% agar | 42°C |
| YPDANISO50 | 2% bactopeptone; 1% yeast extract; 2% glucose; 2% agar; anisomycin 50 µg/ml | 30°C |
| YPDBENOMYL200 | 2% bactopeptone; 1% yeast extract; 2% glucose; 2% agar; benomyl 200 µg/ml | 30°C |
| YPDCHX05 | 2% bactopeptone; 1% yeast extract; 2% glucose; 2% agar; cycloheximide 0.5 µg/ml | 30°C |
| YPDDMSO | 2% bactopeptone; 1% yeast extract; 2% glucose; 2% agar; DMSO 6% | 30°C |
| YPDETOH | 2% bactopeptone; 1% yeast extract; 2% glucose; 2% agar; ethanol 15% | 30°C |
| YPDFLUCONAZOLE | 2% bactopeptone; 1% yeast extract; 2% glucose; 2% agar; fluconazole 20 µg/ml | 30°C |
| YPDFORMAMIDE4 | 2% bactopeptone; 1% yeast extract; 2% glucose; 2% agar; formamide 4% | 30°C |
| YPDFORMAMIDE5 | 2% bactopeptone; 1% yeast extract; 2% glucose; 2% agar; formamide 5% | 30°C |
| YPDHU | 2% bactopeptone; 1% yeast extract; 2% glucose; 2% agar; hydroxyurea 30 mg/ml | 30°C |
| YPDLICL250MM | 2% bactopeptone; 1% yeast extract; 2% glucose; 2% agar; LiCl 250 mM | 30°C |
| YPDMV | 2% bactopeptone; 1% yeast extract; 2% glucose; 2% agar; methylviologen 20 mM | 30°C |
| YPDNACL1M | 2% bactopeptone; 1% yeast extract; 2% glucose; 2% agar; NaCl 1M | 30°C |
| YPDNACL15M | 2% bactopeptone; 1% yeast extract; 2% glucose; 2% agar; NaCl 1.5M | 30°C |
| YPETHANOL | 2% bactopeptone; 1% yeast extract; 2% ethanol; 2% agar | 30°C |
| YPGALACTOSE | 2% bactopeptone; 1% yeast extract; 2% galactose; 2% agar | 30°C |
| YPGLYCEROL | 2% bactopeptone; 1% yeast extract; 2% glycerol; 2% agar | 30°C |
| YPRIBOSE | 2% bactopeptone; 1% yeast extract; 2% ribose; 2% agar | 30°C |
| YPSORBITOL | 2% bactopeptone; 1% yeast extract; 2% sorbitol; 2% agar | 30°C |
| YPXYLOSE | 2% bactopeptone; 1% yeast extract; 2% xylose; 2% agar | 30°C |

Table S.3: Overview of *S. cerevisiae* phenotypes. The phenotypes are calculated as the growth ratio between the listed stress conditions and the standard growing condition for *S. cerevisiae*, namely a medium composed of 2% bactopeptone, 1% yeast extract, 2% glucose, and 2% agar, at 30°C.

| **id** | **univariate GWAS** | **original study [S17]** |
|---|---|---|
| YPACETATE | 15S_rRNA_2 | - |
| YPD14 | - | - |
| YPD40 | - | - |
| YPD42 | Q0045 | - |
| YPDANISO50 | YER180C, YHR164C, YIL147C, YNL205C, YOR009W, YOR127W, YOR138C | - |
| YPDBENOMYL200 | YLR395C | YLR395C |
| YPDCHX05 | - | - |
| YPDDMSO | - | - |
| YPDETOH | - | - |
| YPDFLUCONAZOLE | - | - |
| YPDFORMAMIDE4 | - | - |
| YPDFORMAMIDE5 | - | - |
| YPDHU | YDL056W | YDL056W |
| YPDLICL250MM | YER119C, YGR249W, YML094W, Q0050, Q0065, Q0045 | - |
| YPDMV | - | - |
| YPDNACL1M | YPR083W | YPR083W |
| YPDNACL15M | - | - |
| YPETHANOL | - | - |
| YPGALACTOSE | - | YMR163C, YKR094C - YKR095W; YLR423C - YLR424W |
| YPGLYCEROL | YLR006C, Q0045 | YLR006C |
| YPRIBOSE | - | YLR006C; YIL052C - YIL051C |
| YPSORBITOL | YDR096W, YLR006C | YDR096W, YLR006C |
| YPXYLOSE | - | - |

Table S.4: Overview of results obtained with the univariate GWAS analyses on the *S. cerevisiae* phenotypes. The column named “univariate GWAS” refers to the analysis we performed, while “original study” refers to the single SNP GWAS analysis done by Peter *et al.*. We report the genes onto which the statistically associated SNPs can be positionally mapped. As appreciable from the table, there is consistency for five phenotypes. The witnessed differences may be due to differing SNPs search spaces and type of multiple testing correction used.

| chromosome | SNP | position | gene |
|---|---|---|---|
| 2 | rs1537 | 52083 | YBL088C |
| 2 | rs1555 | 55192 | YBL088C |
| 2 | rs2081 | 121589 | YBL052C |
| 2 | rs3760 | 404662 | YBR081C |
| 4 | rs16176 | 1137328 | YDR334W |
| 4 | rs16184 | 1138235 | YDR334W |
| 4 | rs18032 | 1412942 | YDR477W |
| 5 | rs21237 | 379893 | YER110C |
| 5 | rs21243 | 381575 | YER110C |
| 5 | rs21246 | 381978 | YER110C |
| 7 | rs25594 | 194505 | YGL163C |
| 7 | rs27623 | 480515 | YGL008C |
| 10 | rs38621 | 70181 | YJL194W |
| 10 | rs42210 | 559813 | YJR066W |
| 10 | rs42233 | 562628 | YJR066W |
| 10 | rs42239 | 562775 | YJR066W |
| 10 | rs42885 | 645972 | YJR119C |
| 10 | rs42886 | 646155 | YJR119C |
| 12 | rs55469 | 957510 | YLR418C |
| 12 | rs55977 | 1020763 | YLR442C |
| 13 | rs56319 | 23620 | YML124C |
| 13 | rs58911 | 414299 | YMR075W |
| 14 | rs62671 | 18355 | YNL330C |
| 14 | rs62977 | 48880 | YNL312W |
| 14 | rs64341 | 207220 | YNL236W |
| 14 | rs64691 | 258013 | YNL206C |
| 14 | rs66378 | 461930 | YNL088W |
| 15 | rs71312 | 378902 | YOR025W |
| 15 | rs71489 | 402926 | YOR038C |
| 15 | rs73162 | 616666 | YOR151C |
| 15 | rs75222 | 884597 | YOR304W |
| 16 | rs78517 | 263119 | YPL153C |

Table S.5: Subset of the SNPs identified by *post hoc* Lasso analysis on the SNPs included in the two statistically associated neighborhoods found by networkGWAS ($K_{lin}$) on the natural *S. cerevisiae* phenotype YPGALACTOSE.
